# Torsion is a Dynamic Regulator of DNA Replication Stalling and Reactivation

**DOI:** 10.1101/2024.10.14.618227

**Authors:** Xiaomeng Jia, Xiang Gao, Shuming Zhang, James T. Inman, Yifeng Hong, Anupam Singh, Smita Patel, Michelle D. Wang

## Abstract

The inherent helical structure of DNA dictates that a replisome must rotate relative to DNA during replication, presenting inevitable topological challenges to replication. However, little is known about how the replisome progresses against torsional stress. Here, we developed a label-free, high-resolution, real-time assay to monitor replisome movement under torsion. We visualized the replisome rotation of DNA and determined how the replisome slows down under torsion. We found that while helicase or DNA polymerase (DNAP) individually is a weak torsional motor, the replisome composed of both enzymes is the most powerful DNA torsional motor studied to date. It generates ∼ 22 pN·nm of torque before stalling, twice the stall torque of *E. coli* RNA polymerase. Upon replisome stalling, the specific interaction between helicase and DNAP stabilizes the fork junction; without it, the fork can regress hundreds of base pairs. We also discovered that prolonged torsion-induced stalling inactivates the replisome. Surprisingly, DNAP exchange, mediated by the helicase, is highly effective in facilitating replication restart, but only if excess DNAP is present during stalling. Thus, helicase and DNA polymerase work synergistically as a powerful torsional motor, and their dynamic and fluid interactions are crucial for maintaining fork integrity under torsional stress. This work demonstrates that torsion is a strong regulator of DNA replication stalling and reactivation.

---

Upon the discovery that DNA is a right-handed double helix, Watson and Crick immediately called attention to the formidable topological challenges encountered by DNA replication(*1, 2*). The helical nature of DNA dictates that rotational motion is inherent to replication – for every 10.5 bp (helical pitch) of DNA replicated, the replisome must rotate one turn relative to the parental DNA, leading to over-twisting of the DNA (Fig. 1a; Fig. S1). The resulting (+) DNA supercoiling cannot dissipate fully at the distal ends of the DNA(*3–5*) and must be relaxed by topoisomerases, essential enzymes for topological resolution(*6*). However, even the native full complement of topoisomerases is insufficient to fully relieve torsional stress across the genome(*7–14*). During replication, torsional stress is found especially at regions near replication termination(*15–20*), at DNA fragile sites(*21*), and at conflicts with other motor proteins such as an RNA polymerase(*22, 23*). This indicates that topoisomerases cannot always keep up with the torsional load of genome replication.

**Figure 1.**
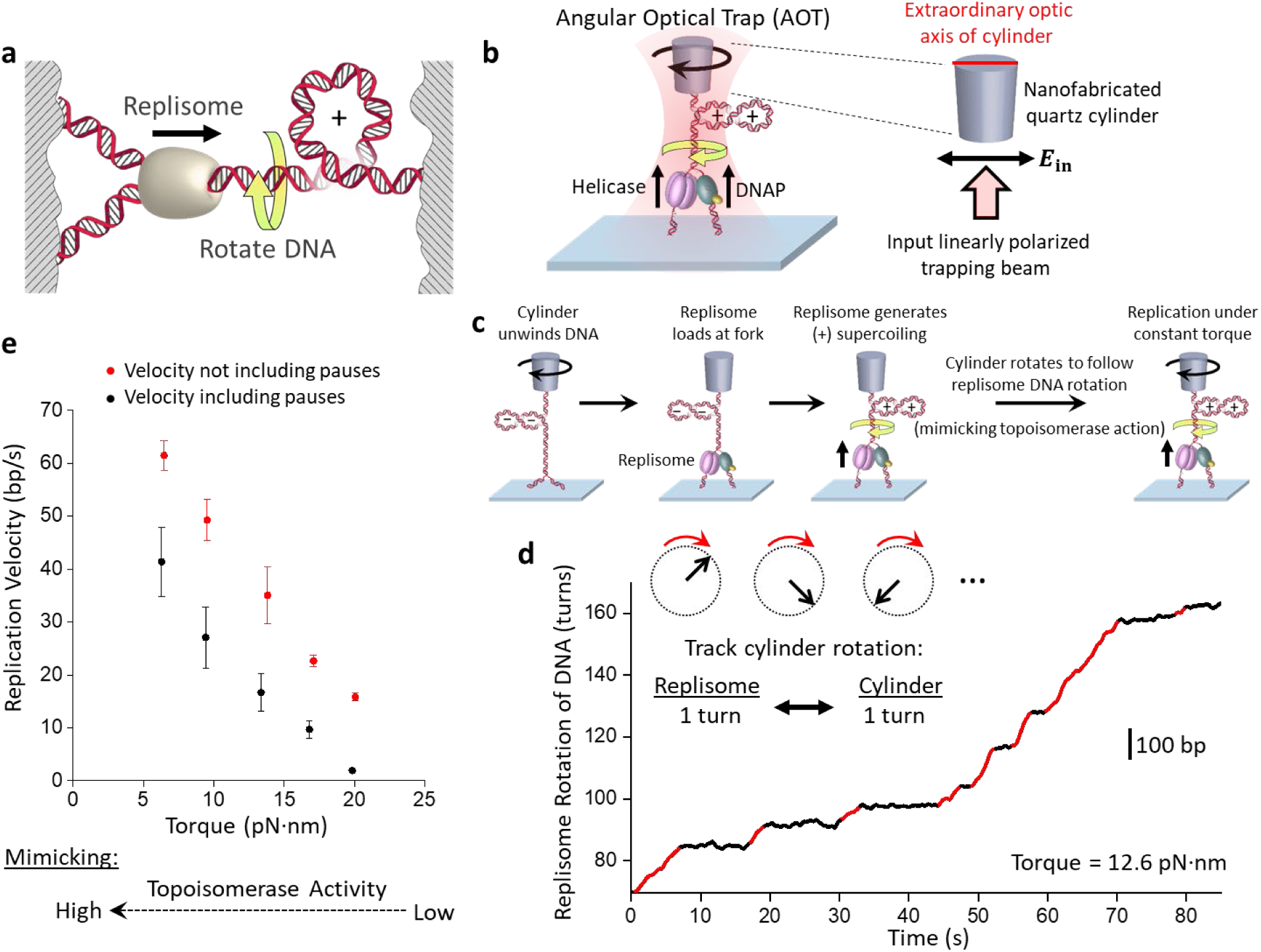
Visualizing replisome rotation of DNA under defined torsion. **a.** A cartoon depicting replisome rotation of DNA during replication. Replication generates extra turns in the DNA substrates. If these extra turns cannot be fully dissipated at the DNA ends or relaxed by topoisomerases, they will produce (+) torsion in DNA. **b.** An experimental configuration to track replisome rotation of DNA using the angular optical trap (AOT) (Methods). A Y-shaped DNA substrate is torsionally constrained between the bottom of an optically trapped quartz cylinder and the surface of a microscope coverslip. Replisome progression generates (+) supercoiling and rotates the cylinder. The replisome rotation is visualized by the cylinder rotation. The extraordinary optical axis of the cylinder tends to align with the polarization of the input trapping beam (right panel) and the cylinder’s angular orientation is precisely detected by the torque detector of the AOT. **c.** Experimental scheme used to track replisome rotation of DNA. The replication fork is underwound to facilitate fork opening and replisome loading by (−) supercoiling the parental DNA. After a T7 replisome loads at the fork and replication starts, the replisome rotates the parental DNA, adding (+) supercoils to the parental DNA, causing it to buckle to form a plectoneme. Once a desired torque is reached, the cylinder is rotated to follow the replisome rotation to prevent further torsion buildup. **d.** An example trace of replisome rotation of DNA under a 12.6 pN·nm torque as visualized via the cylinder (the corresponding Movie is shown in Movie S1). Continuous replication (red regions) is interrupted by pauses (black regions). The scale bar provides the conversion to the translocation distance of the replisome. **e.** Torque-velocity relation of replication. Each data point is obtained from *N* = 14-16 individual traces.

Replication generates torsion, which, in turn, may regulate replication. Unlike stress from local obstacles (such as bound proteins and DNA lesions), torsion acts over distance and can impact regions separated by thousands of base pairs(*24*). Torsion ahead of a replisome may dissociate bound proteins(*25*) or stall oncoming motors(*26, 27*); torsion behind a replisome can entangle daughter DNA strands hindering chromosome segregation(*28, 29*). Importantly, excessive torsion can lead to replication fork stalling, but how individual motors in the replisome impact this process and the role they play in subsequent fork restart and/or DNA damage repair remains enigmatic.

Because of DNA’s inherent helical nature, replication under torsion is a fundamental problem in biology. However, this problem is exceedingly complex, presenting significant challenges for conceptualization and experimentation. There has been little mechanistic understanding of how a replisome elongates against torsion and how this torsion, in turn, impacts replication dynamics; this is mostly due to a lack of direct methodologies for investigation.

## Visualizing replisome rotation of DNA

The fundamental cause for torsional stress during replication stems from the obligatory replisome rotation of the DNA. Understanding the consequences of this rotation requires an experimental method to track the rotational dynamics, which has not been possible for DNA replication. Thus, we developed a method to directly visualize replisome rotation of DNA in real time under a defined torsion. This method was implemented using an angular optical trap (AOT) (Fig. 1b), which is ideally suited to study DNA rotational motion and torsional stress(*26–28, 30–33*). A defining feature of the AOT is its trapping particle. Unlike conventional optical tweezers which typically trap a polystyrene or silica microsphere, an AOT traps a nanofabricated quartz cylinder(*30*) (Fig. S2). The AOT accurately detects the rotational orientation of the cylinder via a change in the laser polarization after laser interaction with the cylinder(*30, 34*) and thus informs the rotational orientation of the attached DNA molecule (Fig. 1b). Here, we have used the cylinder to track the replisome rotation of DNA under a defined torsion (torque).

We enabled this method using the T7 replisome, a model system for studying replication. This minimal system consists of T7 helicase, T7 DNA polymerase (DNAP), and the DNAP processivity factor thioredoxin(*35, 36*) (Fig. 1b; Methods). To begin the experiment, after torsionally anchoring a Y-shaped DNA substrate (Fig. S3; Fig. S4) between the coverslip and a quartz cylinder held in the AOT, we use the AOT to unwind the DNA to facilitate replisome loading at the fork (Fig. 1c). Subsequently, with the cylinder angle held constant, the replisome rotates DNA, converting the (−) supercoiling to (+) supercoiling which then hinders replication, generating increasing (+) torsion as the replisome proceeds. Once a torque value of interest is reached, we allow the cylinder rotation to follow the replisome rotation of DNA, thus limiting further torsion build up and maintaining a constant torque. Therefore, for each turn the replisome rotates the DNA (∼ 10.5 bp), the cylinder also rotates a turn. Because the cylinder’s extraordinary optic axis angle can be tracked at exceedingly high spatial and temporal resolution, this method provides unprecedented resolution of the replication fork dynamics.

Using this method, we directly visualized fork motion in real-time under a specified torque. In the example curve shown in Fig. 1d (Movie S1), we placed the replisome under 12.6 pN·nm of torque to intentionally slow down the replisome for real-time visualization. This rotational trajectory provides detailed information on the replisome motion over a thousand base pairs; the replisome rotates DNA at about 4 turns/s, and the continuous rotation is interrupted by pauses, which reflect a transient halt in the replication motion, likely caused by DNAP entry into the exonuclease state(*37, 38*).

*In vivo*, topoisomerases relax DNA torsional stress. Thus, DNA torsion depends on the extent of topoisomerase activity. Here, we mimicked the extent of topoisomerase activity by specifying a torque value (Fig. 1e). At a low torque (mimicking high topoisomerase activity), the replisome replicates at a high velocity. As the torque increases (mimicking a decrease in topoisomerase activity), the replisome’s pause-free velocity (Methods) decreases with a concurrent increase in pausing, leading to a significantly lower velocity including pauses. The resulting torque-velocity relation specifies how the replisome slows down in response to torsion, a relation characteristic of the chemo-mechanical properties of the torsional motor.

Thus, by directly visualizing replisome rotation of DNA, we demonstrate that the replisome is a torsional motor, capable of rotation against torsional stress. Although this rotational motion is inherent to the helical nature of the DNA, our direct visualization highlights that rotation is an inevitable consequence of replication, paving the way for understanding the impact of torsional stress on fork stability.

## Replisome is a powerful torsional motor

*In vivo*, torsional stress generated by replication increases as the replisome elongates and proceeds to termination(*17, 39–43*), suggesting that topoisomerases are inefficient at regions near replication termination(*44*). In addition, topoisomerase inhibition leads to rapid replisome fork slowdown and stalling in dividing cells(*45–48*), highlighting the crucial need for topoisomerase relaxation of replication-generated torsional stress.

To investigate the dynamics of replisome stalling under torsion, we modified the method depicted in Fig. 1c. Instead of following the replisome rotation of DNA, we restrict the cylinder rotation, thus allowing the DNA substrate to accumulate (+) torsion until the replisome comes to a stall (Fig. 2a). We directly measured the DNA extension, the force, and the torque on the DNA during the process of stalling (Fig. 2b,c) and obtained the fork position from the measured extension (Methods; Fig. S5). Fig. 2b,c show example traces where the replication fork position is monitored as torsion accumulates. The Y-shaped DNA substrate initially does not contain any ssDNA that is required for helicase loading. Consistent with this, we detected no fork activity when helicase alone was present in the reaction buffer, suggesting that DNAP must load onto the template first and replicate a region before subsequent helicase loading. We observed that once both motors load at the fork, the fork moves forward rapidly until it stalls under torsion. We found that the stall torque measured directly using our torque detector agrees well with that of the known DNA buckling torque (Fig. S6), indicating that the replisome was sufficiently powerful to buckle the parental DNA into a plectoneme at a stall.

**Figure 2.**
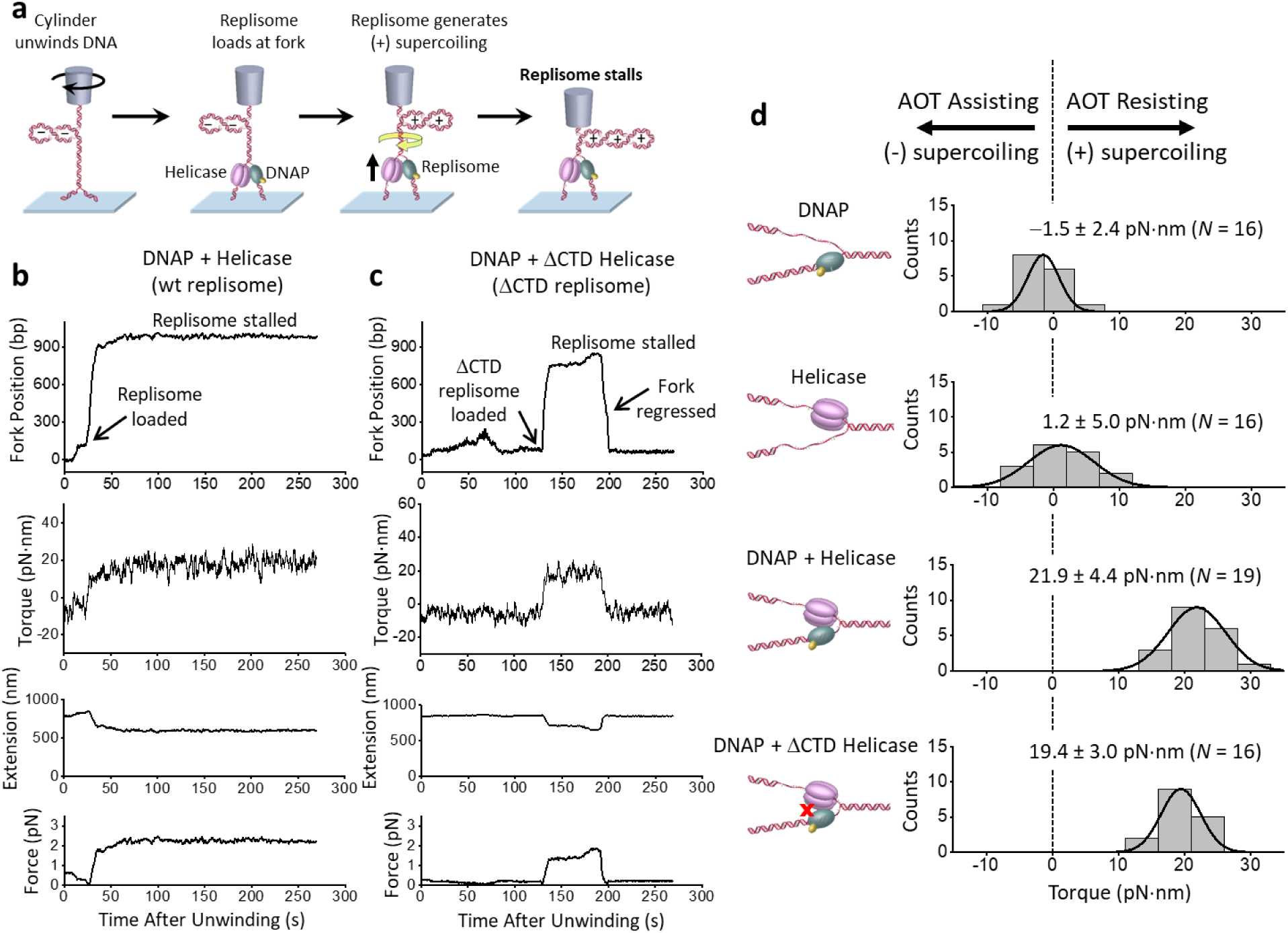
Stalling the replisome under torsion. **a.** Experimental scheme used to stall a replisome under torsion. This experiment starts in the same way as shown in Fig. 1c, but in this case, the cylinder is not allowed to rotate after the initial unwinding. The replication fork is underwound to facilitate fork opening and replisome loading by (−) supercoiling the parental DNA. Upon replisome loading at the fork, the (−) supercoiling is converted to (+) supercoils in the parental DNA, resulting in a decrease in DNA extension and a concurrent increase in force and measured torque. As the (+) torsion generated by replication accumulates, the replisome eventually stalls. **b-c**. Example traces of WT replisome and a replisome containing a ΔCTD helicase. For each trace, force on the DNA, DNA extension, and torque on the DNA are directly measured. The fork position is subsequently obtained from the measured extension (Methods; Fig. S5). Note that the force on DNA is sufficiently small that it does not have any detectable impact on the replication rate (Fig. S7). **d**. Stall torque of the replisome. Shown are histograms of the measured stall torque for DNAP alone, helicase alone, WT replisome, and the ΔCTD replisome. Each histogram is fit by a Gaussian function, and the mean and SD of the Gaussian are indicated. *N* is the number of traces in each histogram. A negative torque indicates that an enzyme requires an assisting torque to enable forward motion, whereas a positive torque indicates that an enzyme can move forward against a (+) torsion.

A replisome converts chemical energy to perform the mechanical work of rotating DNA. This mechanical work can then be used to move forward against the torsional stress, dissociate bound proteins, and reconfigure DNA structures and topology(*49, 50*). Thus, we directly measured the torque to stall a wild-type (WT) replisome (Fig. 2b), since the stall torque is an inherent property of a torsional motor. We found the WT replisome stalls at a torque of 21.9 ± 4.4 pN·nm (mean ± SD), making it a powerful torsional motor (Fig. 2d). To put this value in perspective, *E. coli* RNA polymerase (RNAP) on its own can generate ∼ 11 pN·nm of torque(*26*), which is enough to melt DNA(*32, 33*), and a torque of +19 pN·nm has been shown to significantly facilitate the dissociation of histone H2A/H2B from a nucleosome(*25*). The T7 replisome is therefore about twice as powerful in terms of torque generation capacity and is the most powerful DNA torsional motor studied to date.

It is possible that the T7 replisome torsional capacity simply results from the additive torsional capacities of the two motors at the fork. To investigate this possibility, we measured the stall torque of DNAP alone without helicase and found it to be –1.5 ± 2.4 pN·nm (Fig. 2d), revealing that DNAP alone has minimal capacity to generate (+) torsion despite bearing some structural and functional similarities to RNAP. To determine the stall torque of the helicase alone, we used a modified Y-shaped DNA substrate containing a ssDNA region for helicase loading (Methods). We found the stall torque to be 1.2 ± 5.0 pN·nm (Fig. 2d), showing that helicase alone also has minimal capacity to generate (+) torsion.

We wondered if the replisome’s high torsional capacity is a result of the specific interaction between the C-terminal domain (CTD) of the helicase and DNAP, as this interaction is essential for T7 phage replication(*51*). We thus carried out a similar stalling experiment with a replisome containing a ΔCTD helicase (referred to as ΔCTD replisome) (Fig. 2c). The ΔCTD replisome also generates significant torsion, albeit being less stable upon stalling, with the fork position sometimes undergoing significant reverse movement (Fig. 2c). Surprisingly, we found that a replisome containing a ΔCTD helicase (referred to as ΔCTD replisome) remains a powerful torsional motor, generating a torque of 19.4 ± 3.0 pN·nm (Fig. 2d), only marginally smaller than that of the wild-type (WT) replisome.

Therefore, these results indicate that the replisome’s torsional capacity is primarily due to the presence of the two motors at the fork. Each motor in the replisome on its own does not afford any strong capacity to work against torsion, likely because helicase can slip under the influence of the fork(*52*) and DNAP can also reverse translocate(*53*). However, working in conjunction, the two motors at the fork convert the replisome into a remarkably powerful torsional motor. This synergistic cooperation does not rely on any known specific interactions between the two motors but may be a result of one motor keeping the fork open for the other motor by limiting reverse fork motion(*54–56*) and augmenting DNA breathing at the fork(*57*). Unlike RNAP that holds both DNA strands within the motor, the replisome splits the two strands, with each motor tracking one of the strands(*36*). This configuration can create a larger lever arm for torque generation around the parental DNA’s central axis, akin to using a corkscrew with a longer handle to screw into the cork of a wine bottle.

## Torsion leads to fork regression

We found that once the replisome comes to a stall, the fork is not static and instead exhibits continuous forward and reverse movement (Fig. 2b,c). To characterize fork stability during a stall, we aligned the fork position of each trace at its maximum value and plotted how the fork position regresses over time, leading to a concurrent torsion reduction (Fig. 3a). We found the fork of the WT replisome is comparatively stable, with the fork regressing about 80 bp over a minute. In contrast, the ΔCTD replisome undergoes a more dramatic fork regression of 240 bp in a minute, with a larger concurrent reduction in torsion, reflecting a reduced ability to sustain torsional stress. Interestingly, although the ΔCTD replisome can still generate a high stall torque (Fig. 2d), this mutation significantly impairs fork stability during a stall.

**Figure 3.**
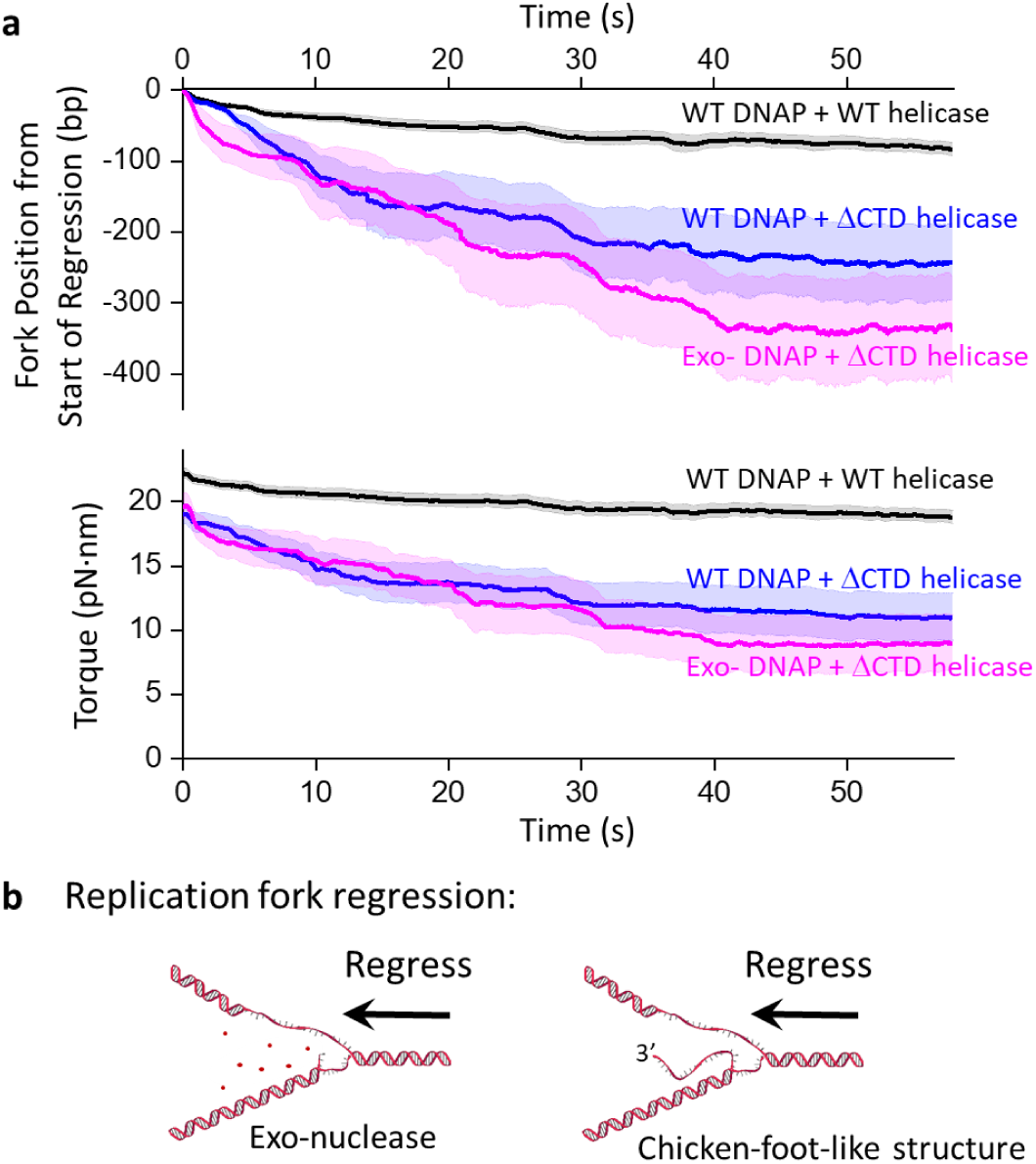
Fork regression dynamics during stalling under torsion. **a.** Fork regression versus time. All traces are aligned in position and time at their maximum fork position. For each type of replisome, we show the mean fork position versus time, with their SEM bracketing the shaded regions. The total number of traces are: 48 (wt replisome), 22 (ΔCTD replisome), and 12 (ΔCTD exo-replisome). **b.** Cartoons illustrating two possible mechanisms of fork regression. Enzymes bound near the fork are omitted for clarity of the fork configuration.

The observed fork regression can result from two distinct mechanisms. Since DNAP has an exonuclease activity, the replisome may reverse translocate by DNAP removing the replicated nucleotides (Fig. 3b left panel). Alternatively, the torsional stress generated by the replisome could reverse the fork by squeezing the replicated strand off the template DNA, forming a structure that resembles a chicken-foot (Fig. 3b right panel). Previous studies show that replication stress can ultimately lead to the formation of a chicken-foot structure(*15, 45, 58*). Because such a structure maintains the same number of base pairings in the DNA structure during the regression, this type of fork reversal is likely energetically mobile and can proceed over a longer distance while releasing torsional stress.

Here, we investigate the formation of a “chicken-foot-like” structure using a minimal system composed of only the replisome components. We stalled the replisome containing a ΔCTD helicase and an exo-DNAP (referred to as ΔCTD exo-replisome; Methods), which lacks exonuclease activity and can only form a chicken-foot-like structure under fork regression. We found that the ΔCTD exo-replisome also displays long-distance fork regression (reversing about 340 bp in a minute), similar to but at a slightly greater extent than the ΔCTD replisome, suggesting that the fork regression observed for the ΔCTD replisome may primarily be a result of the chicken-foot-like structure formation. These findings demonstrate the crucial role of the specific interaction between the helicase and DNAP in stabilizing the replication fork under torsional stress by limiting fork reversal. In the absence of this interaction, the fork may collapse via the formation of chicken-foot-like structures, rendering the replisome inactive.

Our experiments show that the replisome is dynamic when stalled under torsion, and even the WT replisome undergoes moderate fork regression. Since fork regression releases torsional stress, such a regression could be a protective mechanism to tolerate the torsional stress and prevent genome instability(*59–62*). Our work examines how fork regression proceeds when only a minimal replication machinery is present. *In vivo*, fork reversal under stress is further modulated by other enzymes, such as annealing DNA helicases, which regulate fork reversal to safeguard genome integrity(*59, 63–65*).

## Prolonged torsion limits fork restart

*In vivo*, if the replisome is stalled by torsion, torsion may be relaxed by the arrival of topoisomerases. Subsequent replication restart is crucial for maintaining genomic integrity and preventing mutations and is regulated by a multitude of enzymes and factors(*61, 66*). Here, we investigated what governs replication restart in the presence of the minimal replication machinery proteins. For these experiments, we allowed the replisome to stall under torsion for a specified time, mimicked topoisomerase torsional relaxation by unwinding DNA to reduce the torsion, and examined the replisome’s ability to restart replication (Fig. 4a).

**Figure 4.**
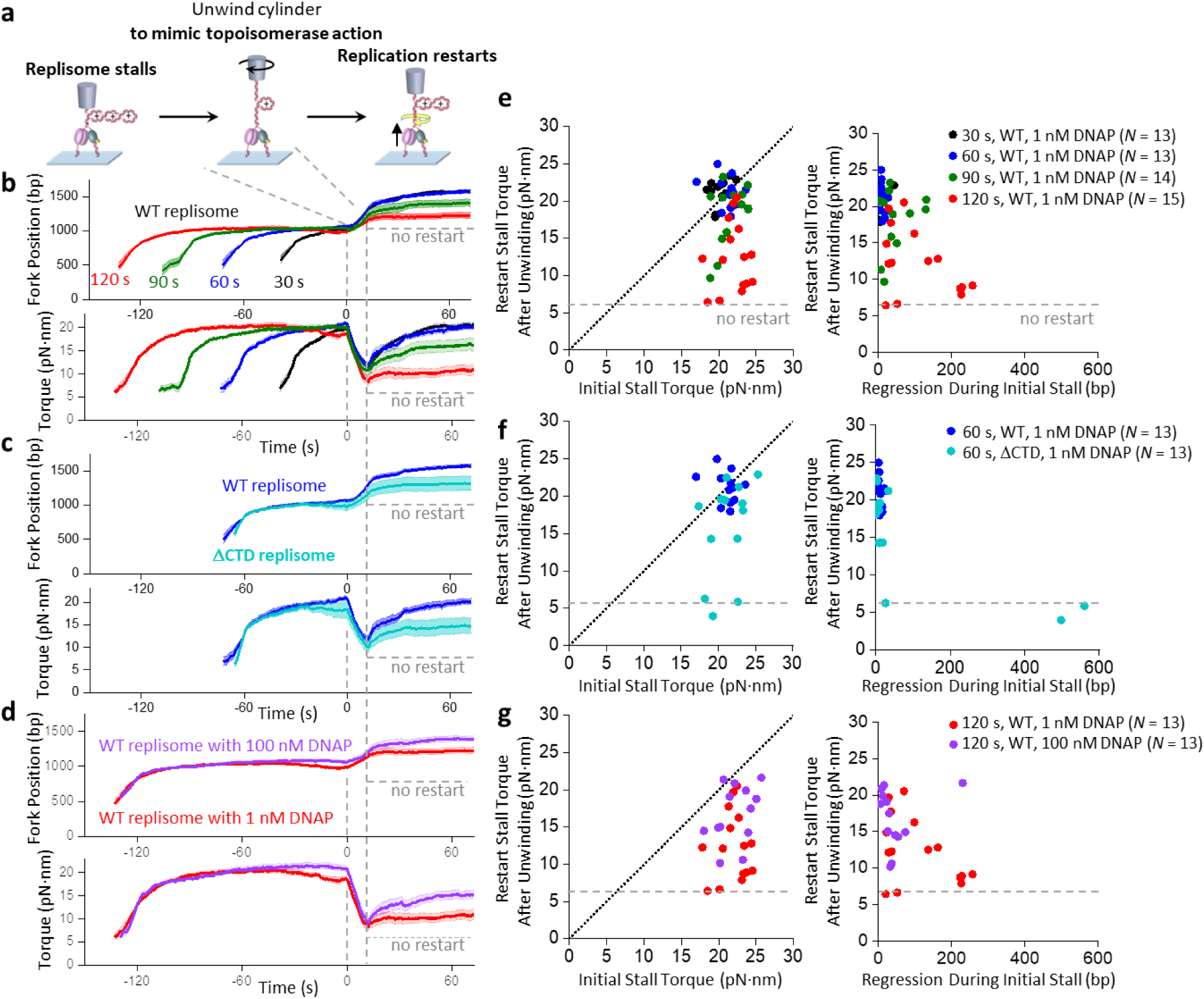
Replication restart after stalling. **a.** Experimental scheme for replication restart using the angular optical trap. The replisome is first loaded at the fork and allowed to accumulate (+) supercoiling and stall (as shown in Fig. 2). Subsequently, we unwind the cylinder to release some of the torsional stress, mimicking the action of topoisomerases. After unwinding, replication may resume, leading to the accumulation of (+) supercoiling and stalling of the replisome. **b**-**d**. Replication restart under different stalling times and protein compositions. Shown are the mean curves after aligning all traces at the start of the AOT unwinding. For each condition, the mean curve of *N* = 12-16 traces is bracketed by with the SEM (shaded region). **e-g**. Replication restart characterization from traces used in **b-d**, respectively. The effectiveness of restart is characterized by the restart stall torque after the unwinding step. The diagonal dashed lines represent when the restart stall torque is the same as the initial stall torque, indicating an effective restart. Each data point came from a single trace. The three panels on the right show how the ability to restart correlates with the regression distance during the stalling.

We first examined a condition of partial torsional relaxation, which could occur *in vivo* when topoisomerases cannot remove all torsion generated by replication (Fig. 4b-d). In the torsion relaxation step of the experiment, we used the AOT cylinder to unwind DNA to reduce the torsion to +6 pN·nm, which is well below the stall torque and requires unwinding of no more than 50 turns if the replisome is inactive during the relaxation. However, in cases where the replisome becomes immediately active during torsion relaxation and generates a torque greater than +6 pN·nm, we unwound 50 turns. After torsion relaxation, if the replisome is fully active, it should restart replication, generate torsion, and stall again under torsion. We use the restart stall torque after the unwinding step as a measure of restart efficiency; a fully active replisome after the restart should reach a torque close to the initial stall torque.

We found that when the replisome is initially stalled for 30 s or 60 s, replication restarts efficiently upon torsion reduction before stalling again at a similar stall torque (Fig. 4b,e). However, as the initial stall time increases to 90 s and 120 s, the replication undergoes more significant fork regression during the initial stall, with the restart stall torque reduced from the initial stall torque (Fig. 4b,e). The results demonstrate that prolonged stalling inactivates replication, highlighting the importance of timely torsional relaxation by topoisomerases *in vivo*. A delay in topoisomerase arrival can lead to prolonged replication stalling under torsion, rendering the fork less active.

We then examined the role of DNAP-helicase interaction in the replication restart (Fig. 4c,f). We found the ΔCTD replisome reaches an initial stall torque similar to that of the WT replisome but undergoes more long-distance fork regression during the stall and has a reduced restart stall torque. Thus, the specific interaction between DNAP and helicase not only stabilizes the fork during stalling but also promotes replication restart after stalling.

Importantly, we were particularly intrigued by the low restart rate under the 120 s stalling time (Fig. 4b,g), as the inability to restart replication is detrimental to genome stability. If the low restart rate results from DNAP dissociation under stress, then DNAP must reload before replication restarts, and inefficient DNAP reloading could limit the restart. However, this is unlikely since the DNAP loading time on the initial DNA substrate is short under our experimental conditions (Fig. S8), suggesting an increase in DNAP concentration should not increase the restart rate. Surprisingly, when we increased the DNAP concentration from 1 nM to 100 nM, the replisome restarted more efficiently (Fig. 4d,g). Under 100 nM DNAP, the replisome also remains more active even during the stall, showing less fork regression and generating a slightly higher torque. Thus, the excessive DNAP stabilizes the fork during a stall, facilitating subsequent replisome restart. This finding also raises many additional questions. Is the low restart rate after a long stall due to incomplete torsion relaxation? Would excessive helicase facilitate restart when the DNAP loading rate is not limiting? Does an excessive amount of other proteins play a similar role in the restart?

## DNAP exchange promotes fork restart

Answering these questions required many additional experiments under different conditions. The scope of these experiments motivated us to develop a new method based on magnetic tweezers (MT)(*28, 67, 68*), enables parallel measurements of multiple replisomes under torsion. The experimental procedure for replication restart using the MT (Fig. 5a) is similar to that using the AOT (Fig. 4a). Although this MT method does not allow direct torque measurements, it can readily monitor replication elongation, stalling, and restart, while the torque can still be obtained indirectly from the force since stalling occurs when the DNA is buckled to form a plectoneme according to our AOT data (Fig. S6). Under the 1.0 pN force used in these assays, (+) DNA buckling occurs at 12.6 pN·nm torque, which could only be generated by an active replisome (Fig. 2d). Thus, we classify a trace as having restarted if the replisome can (+) buckle the DNA after the unwinding step (Fig. 5b).

**Figure 5.**
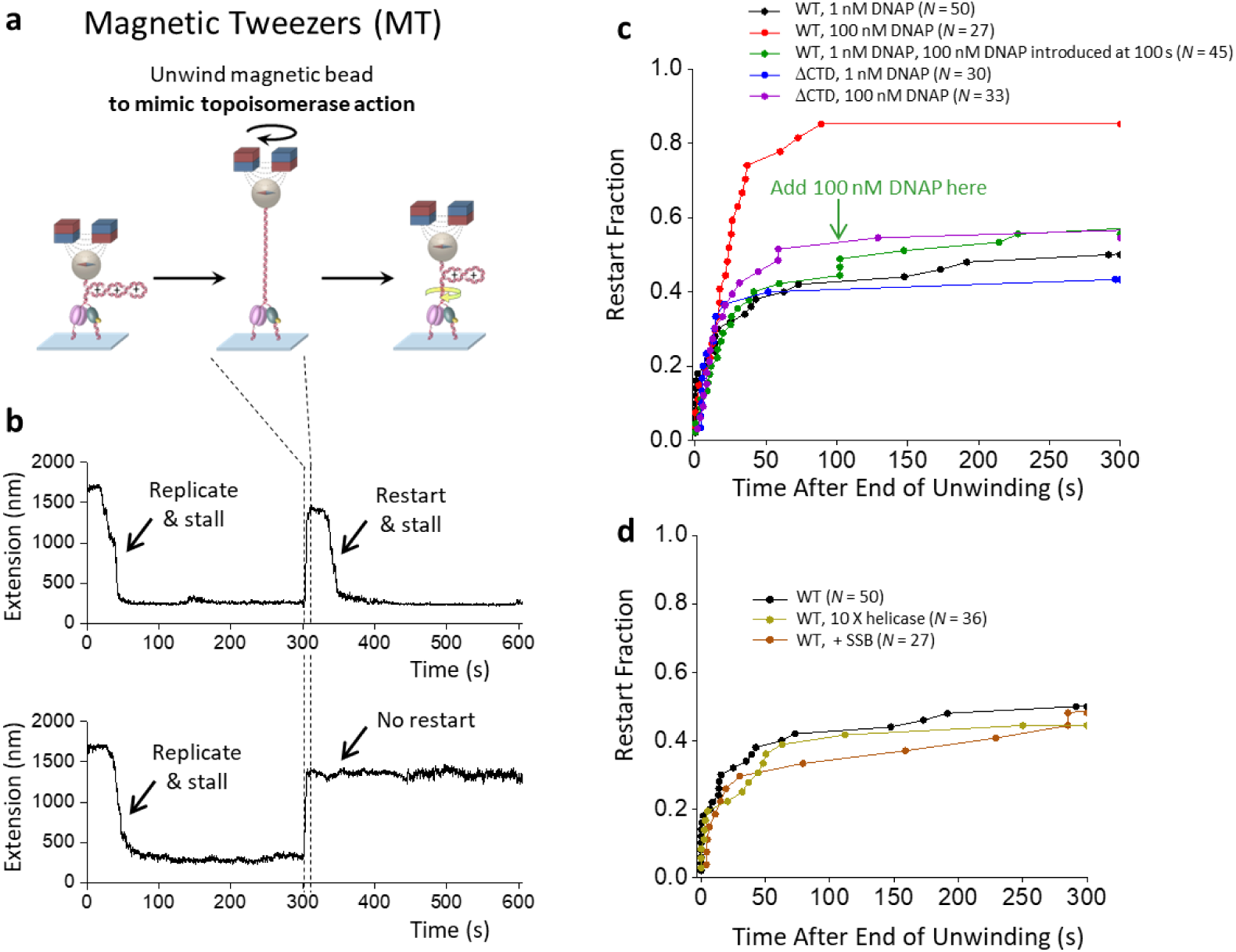
DNA exchange during replication restart. **a.** Experimental scheme for replication restart using magnetic tweezers. As with the AOT experiments in Fig. 4, the replisome is first allowed to load and accumulate (+) supercoiling and stall. Subsequently, we unwind the magnetic bead to release all torsional stress, mimicking effective torsional relaxation by topoisomerases. After unwinding, replication may resume, leading to the accumulation of (+) supercoiling and stalling of the replisome. **b.** Two example traces. DNA tethers are held under 1.0 pN. In each trace, the initial replication leads to the generation of (+) torsion, as evidenced by an extension decrease (also see Fig. 4). Continued replication should eventually stall the replisome. The subsequent unwinding of the DNA fully relaxes the (+) torsion, increasing the DNA extension. Following this unwinding step, the ability of the replisome to restart is examined. The top trace shows an example of replisome being able to restart, as revealed by the extension decrease. The bottom trace shows an example of the replisome being unable to restart, where the extension remains essentially unchanged. **c.** The effect of DNAP concentration on replication restart efficiency. The green arrow indicates the time when we introduced 100 nM DNAP together with helicase into the reaction for the green curve. **d.** The effect of helicase concentration and the presence of SSB on replication restart efficiency. The olive curve was obtained by increasing the helicase concentration to 10 X of that used in the standard condition (Methods). The maroon curve was obtained by including 200 nM SSB in the standard condition.

Using the MT method, we focused on factors that can facilitate replication restart after a long stall (240 ± 17 s; mean ± SD). To fully eliminate torsion, we unwound 100 turns during the torsion relaxation step. This condition should remove any torsional stress and promote the reversal of the fork regression (Methods). This mimics an *in vivo* situation where a replisome is stalled for a long time but subsequent arrival of topoisomerases is highly effective at relaxing torsional stress. We first re-examined the role of excess DNAP in replication restart (Fig. 5c).

We found a marked increase in the restart rate from 40% to 85% when the DNAP concentration increased from 1 nM to 100 nM, reinforcing the observations made using the AOT. Examination of the DNAP loading time again shows that DNAP loading should be rapid even under 1 nM DNAP (∼ 14 s) (Fig. S9). Significantly, this enhanced restart requires excessive DNAP to be present during the initial stall, as increasing DNAP concentration to 100 nM after the prolonged stall is much less effective in promoting replication restart (Fig. 5c, green curve). Thus, the findings using the AOT and MT collectively demonstrate that prolonged replisome stalling by torsion inactivates replication, which can be effectively circumvented by DNAP exchange.

To test this hypothesis further, we examined the role of the helicase CTD, which has previously been shown to mediate the DNAP exchange(*69, 70*) (Fig. 5c, purple curve). Since the T7 helicase is a hexamer, multiple DNAP enzymes can be recruited to the helicase via its CTD, greatly increasing the effective local concentration of DNAP at the fork (on the order of mM) to facilitate the exchange. Indeed, the ΔCTD replisome, lacking the helicase CTD and unable to recruit DNAP, shows minimal increase in the replication restart rate when the DNAP concentration is increased from 1 nM to 100 nM, indicating a crucial role of the helicase CTD in enabling DNAP exchange to promote fork restart. This finding lends strong support for the proposed DNAP exchange mechanism to maintain an active fork.

To determine if the exchange of other proteins, such as helicase and *E. coli* SSB, involved in replication can play a similar role as DNAP exchange in replication restart, we increased the helicase concentration by 10 times from 180 nM to 1.8 µM and found the restart efficiency to be minimally affected (Fig. 5d). SSB has been implicated in facilitating fork progression and might play a role in the restart efficiency(*71*). We found that adding 200 nM SSB in the reaction does not promote replication restart (Fig. 5d).

Thus, our results support a highly significant role of DNAP exchange in maintaining an active fork during the stall and facilitating subsequent replication restart after torsional relaxation. A stalled replisome can remain active if its DNAP can be dynamically replaced during stalling, likely by limiting the formation of DNAP binding to misconfigured forks (Fig. S10). Previous work found DNAP exchange could occur under no replication stress and proposed the intriguing possibility of DNAP exchange as a mechanism of enabling replication to overcome replication stress for subsequent replication restart(*70*). Direct experimental evidence to support this mechanism has been lacking until this study. To our knowledge, our findings provide the first direct experimental demonstration of the crucial role of DNAP exchange in safeguarding the replication fork and facilitating replication restart under torsional stress.

## Discussion

To understand how a replisome responds to torsional stress, we have investigated the T7 replisome containing the minimal replication components. Using this system, we provide unprecedented information on replication dynamics, demonstrating that torsion is a strong regulator of replication. We observe that the fork must be highly dynamic to remain active, and these dynamic behaviors are exhibited in the fork’s response to torsion and interactions among the replisome constituents.

We found the T7 replisome to be a powerful torsional motor (Fig. 2d). The torsional capacity of the replisome is particularly relevant in hard-to-replicate regions. In particular, studies on replication-transcription conflicts demonstrate that head-on conflicts have more dire consequences than co-directional ones, such as replication slow-down, fork stalling, and replisome disassembly that necessitates replication restart(*23, 72*). During a head-on conflict, the replisome and RNAP generate (+) supercoiling ahead, leading to (+) torsion accumulation between the two motors(*73*) (Fig. S11). Studies support a model in which this (+) torsion is the culprit for replication stress during the head-on conflicts(*22, 24, 73*). In addition, a replisome ultimately wins in this head-on conflict(*74*), but the mechanism remains unclear.

Our studies show that the T7 replisome can generate 22 pN·nm of torque (Fig. 2d), twice that of *E. coli* RNAP(*26*). Thus, *in vivo*, the (+) torsion between the two machineries during a head-on conflict could stall and destabilize the bound RNAP first while permitting the replisome to continue to move forward. This could occur well before any physical encounter of the replisome with RNAP, as torsion in the DNA acts over distance. The differential torsional capacities of the two machineries may play a role in how a replisome wins the conflict. Even so, the accumulated (+) torsion may slow down and eventually stall the replisome, as we observed in this work (Fig. 1,2). We found that prolonged stalling under torsion leads to reverse fork motion (Fig. 3) and inactivates the replisome (Fig. 4) which requires alternative mechanisms for restart. These findings highlight the severe consequences of head-on conflicts and the crucial role of topoisomerases in timely torsion relaxation.

We show that an active fork requires dynamic and fluid interactions between the helicase and DNAP. We previously found that the T7 helicase on its own can readily slip backwards(*52*) but becomes highly processive when complexed with a non-replicating DNAP, and this interaction, mediated via the CTD of the helicase, was sufficiently strong to allow helicase to drag the non-replicating DNAP along the DNA(*75*). The current work shows that this interaction also stabilizes the fork, prevents long-distance fork reversal during stalling (Fig. 3a), and facilitates subsequent fork restart (Fig. 4c,f). While this interaction ensures fork stability, it may also present a potential problem for replication restart if the DNAP in the replisome becomes inactive during stalling.

We made a surprising discovery that the T7 replisome resolves this potential issue by DNAP exchange during stalling (Fig. 4d,g; Fig. 5c; Fig. S10). Because the T7 helicase is a homo-hexamer, it can recruit multiple DNAP molecules to the fork(*36, 76*), although only one is engaged in the leading-strand replication. The additional DNAPs are poised to replace the leading-strand DNAP. We found that instead of simply waiting for the DNAP to dissociate, these helicase-bound DNAPs promote its dissociation and subsequently replace it to maintain an active fork. Importantly, we found that replication restart is highly efficient only if excess DNAPs are present during stalling, indicating steady DNAP exchange is necessary to maintain an active fork. This exchange is particularly crucial for prolonged stalls under torsion, which can lead to fork arrest. These findings extend the previous observations of DNAP exchange *in vitro* and *in vivo*(*69, 70, 76–83*) and provide direct support for a model of active replacement via promoting DNAP dissociation(*70, 84*).

Although our work focuses on the T7 replisome, the experimental approaches established here can be broadly applied to other replication systems. While the helicases in the T7, the T4, and the bacterial replisomes are located on the lagging strand, CMGs in eukaryotic replisomes are positioned on the leading strand ahead of the DNAP(*85*). Despite these differences, these replisomes all possess specific interactions between the helicase and the replicative DNAP. These replisomes may differentially regulate their torsional generation capacity, fork regression under torsion, and the subsequent fork restart upon torsional relaxation. This work creates many new opportunities to investigate replication dynamics under torsion.

## ACKNOWLEDGEMENTS

We thank members of the Wang Laboratory for helpful discussion and comments. We especially thank Dr. P.M. Hall for helpful advice on DNA substrate design. This work is supported by the National Institutes of Health grants R01GM136894 (to M.D.W.). M.D.W. is a Howard Hughes Medical Institute investigator. This work has been performed in part at the Cornell NanoScale Facility, a member of the National Nanotechnology Coordinated Infrastructure (NNCI), which is supported by the National Science Foundation (Grant NNCI-2025233).

## AUTHOR CONTRIBUTIONS

X.J., J.T.I., X.G., and M.D.W. designed single molecule assays. X.J. prepared DNA substrates. A.S. and S.S.P. purified and characterized T7 gp4A’, T7 gp4A’ ΔCt, T7 gp5 Exo-, and *E. coli* SSB proteins. X.G. and J.T.I. updated the AOT for replication assays. X.J., X.G., and S.Z. performed single-molecule experiments. Y.H. fabricated quartz cylinders. Y.H., X.G., and X.J. calibrated the cylinders. X.J. and J.T.I. analyzed data. M.D.W. wrote the initial draft. All authors contributed to manuscript revision. M.D.W. supervised the project.

## COMPETING INTERESTS

The authors declare no competing financial interests.

### Code availability

Data analysis routines used to process and generate plots are available in Github: https://github.com/WangLabCornell/replication/

## DATA AND MATERIAL AVAILABILITY

All data are available in the main text or the supplementary materials.

## SUPPLEMENTARY MATERIALS

Materials and Methods

Figs. S1 to S11

Movie S1

