## Supplementary Materials for "Torsion is a Dynamic Regulator of DNA Replication Stalling and Reactivation"

### Materials and Methods

#### Proteins

The M64L mutant of T7 helicase gp4 (T7 gp4A') and its C-tail deletion mutant with 17 C-terminal amino acid residues deleted (gp4A' ΔCt) were purified from *E. coli(1, 2)*. Introduction of M64L mutation inactivates a second start codon in T7 gp4 ORF and prevents expression of gp4B. *E. coli* cells expressing these proteins were lysed by freeze-thaw cycles in lysis buffer (50 mM Tris-HCl, pH 7.5, 100 mM NaCl, 1 mM EDTA, 1 mM DTT, 10% sucrose, 1 mM PMSF and 0.4 mg/ml lysozyme). Polyethylenimine (PEI) was added to the lysis supernatant followed by ammonium sulphate precipitation. The precipitates were solubilized and further purified on phosphocellulose (P-11 resin) and DEAE sephacel chromatography columns.

Exonuclease deficient mutant of T7 gp5 DNA polymerase (D5A, E7A) was also purified(*3*). Briefly, *E. coli* cells expressing T7 gp5 Exo- were lysed in buffer consisting of 50 mM Tris-HCl, pH 8.1, 2.5 mM EDTA, 1 mM β-mercaptoethanol, 0.1 mM DTT, 0.3 mg/mL Lysozyme, 150 mM NaCl and 0.1% sodium deoxycholate. NaCl concentration was gradually increased to 0.5 M while gently stirring the lysate on ice. Lysate was cleared by centrifugation and the protein was precipitated using 0.5% final concentration of PEI, pH8. Protein was purified by ammonium sulfate precipitation. Precipitates were dissolved, dialyzed to remove excess salt and further purified on phosphocellulose (P-11 resin) and DEAE sephacel chromatography columns.

*E. coli* SSB was purified from *E. coli*(*4*). Briefly, *E. coli* cells expressing SSB were lysed, and the lysate was cleared by centrifugation. PEI was added to the cleared lysate to precipitate the protein. Precipitants were extracted in buffer containing 0.4 M NaCl. Protein was further purified by ammonium sulfate precipitation followed by Q-sepharose column chromatography.

Wild-type T7 gp5 DNA polymerase was purchased from New England Biolabs (NEB, Ipswich, MA). *E. coli* thioredoxin was purchased from Sigma-Aldrich (St Louis, MO).

#### DNA substrates

We conducted the majority of experiments using a Y-shaped DNA substrate (Fig. S3), which is composed of a pre-formed replication fork with a pair of ~2.2 kb symmetric DNA strands downstream of the fork and ~5 kb of parental DNA. Each end of the Y-shaped template is ligated to a ~500 bp multi-labeled tethering adaptor(*5-7*). The upstream DNA is synthesized via PCR from pLB601 and the parental DNA from pRL574. The pre-formed replication fork was made of 4 ssDNA oligos (IDT) which are annealed and ligated to upstream and parental DNA(*8*). The pre-formed replication fork DNA sequence does not allow for fork regression past the initial fork position. Finally, the upstream DNA strands are nicked to ensure the daughter strands are torsionally relaxed.

For the helicase only experiments, we used a Y-shaped DNA substrate that is nearly identical to the template described above except containing a small region of ssDNA on the lagging strand near the fork. The only difference is that the pre-formed replication fork was made of 3, rather than 4, ssDNA oligos (IDT). After the 3 ssDNA oligos are annealed, a 27-nt ssDNA region is naturally formed at the fork for helicase loading.

**Experimental conditions**

Single molecule tethers were formed in a nitrocellulose coated sample chamber(*5-7, 9*). The sample chamber surface was functionalized with 25 ng/µL antidigoxigenin (Vector Labs MB-7000) and then passivated with 25 mg/mL β-casein (MilliporeSigma C6905). DNA replication substrates were then incubated at a concentration of 5 pM. Quartz cylinders or magnetic beads (Dynabeads MyOne Streptavidin T1, ThermoFisher 65601) were introduced. The chamber was washed with the replication buffer (50 mM Tris-HCl (pH 7.5), 40 mM NaCl, 1.5 mM EDTA, 10% glycerol, 8mM $MgCl_2$, 1 mM dNTPs, 2 mM DTT, 0.5 mg/mL β-casein) before flowing in the protein of interest. All experiments were performed in the replication buffer.

Unless stated otherwise, replication was conducted under the following conditions: (1) wt replisome with 1 nM wt T7 gp5 DNA polymerase, 180 nM wt T7 gp4A' helicase (monomer), and 100 nM thioredoxin; (2) ΔCt replisome with 1 nM wt T7 gp5 DNA polymerase, 45 nM gp4A' ΔCt helicase (monomer), and 100 nM thioredoxin ; (3) ΔCt exo- replisome with 1 nM exo- gp5 DNA polymerase, 45 nM gp4A' ΔCt helicase (monomer), and 100 nM thioredoxin .

We found that ΔCt helicase can bind to DNA 4X faster than wt helicase, as determined using our standard helicase unwinding assay(*10, 11*). We thus decreased ΔCt helicase concentration by 4X from the wt helicase concentration. In experiments that require 100 nM wt T7 gp5 DNA polymerase, thioredoxin concentration was raised to 1000 nM.

**Single-molecule methods**

Our lab previously developed the angular optical trap (AOT) for simultaneous measurements of force, extension, torque, and rotation of DNA(*12-15*). A defining feature of the AOT is its trapping particle, a nanofabricated quartz cylinder (Fig. 1b; Fig. S2). The angular orientation of the cylinder is controlled by the trapping laser polarization while torque on the cylinder is directly measured by the change in the angular momentum of the laser after interaction with the cylinder. At the beginning of each experiment, the trap center of the AOT was set to be ~ 800 nm above the surface of the coverglass. The tether was unwound by 40 turns which (-) supercoils the DNA and applies (-) torsion to the fork. This (-) torsion weakens the base pairing interactions of the parental DNA at the fork and replisome loading. Once the replisome loads at the fork, replication proceeds rapidly, generating (+) supercoiling in the parental DNA. The stall torque for each trace is defined as the maximum torque measured within 60 s after the start of stalling.

Magnetic tweezers (MT) experiments were performed on our custom-built instrument(*5, 7, 9*). In this instrument, a pair of magnets apply a constant force to a field of DNA tethers via magnetic beads, and rotation of the magnets rotates the beads to introduce supercoiling to the DNA molecules. During a replication restart experiment (Fig. 5c-d), the force on a Y-shaped DNA substrate is kept at 1.0 pN. After the DNA tethers are unwound by 30 turns, replication start is evidenced by an extension decrease, consistent with (+) supercoil accumulation. At 1.0 pN force, when the DNA is unwound, the extension remains rather flat as the DNA undergoes the melting transition(*6, 16*) (Fig. S4); whereas when the DNA is overwound, the extension decreases linearly once the DNA is buckled to form a plectoneme(*6, 17*), and the buckling torque is 12.6 pN·nm (Fig. S4d), which is greater than the torque that DNAP alone or helicase alone can generate and can only be generated by an active replisome.

This break of symmetry provides a convenient way to identify an active replisome by an extension decrease. Continued replication eventually stalls the replisome shortly after the magnetic bead contacts the surface. The extra turns added by replication upon stalling is estimated to be 51 turns based on the stall torque of 22 pN·nm using the torsional stiffness of the buckled DNA and plectonemic DNA(*6*). We then wait for a specified amount of time before unwinding DNA at 10 turn/s by 100 turns, which should remove all (+) turns added by replication, resulting in an increase in the DNA extension. Subsequent replication restart is evidenced by an extension decrease.

**Replication pause analysis**

The replication velocity at a constant torque is obtained from the replisome position versus time plots such as shown Fig. 1d. Replisome activity is monitored for 60 s or until the fork regresses at least 100 bp. The velocity including pauses is a linear fit to the rate of replication during this period. To detect pauses, the position versus time plot after being smoothed by 0.5 s Gaussian filter is used to generate the dwell time histogram versus position(*18-20*). A pause is detected and removed when the dwell time falls below 0.1 s/bp. The pause-free velocity of each trace is a duration weighted mean of the linear fits to position versus time of regions of activity between pauses.

**Replication fork position determination**

The location of the fork is determined from the extension of the DNA tether (Fig. S5). Prior to each experiment, the DNA extension is measured as a function of applied turns by the cylinder. This calibration curve is then used to convert the subsequent extension to the number of turns that have accumulated in the parental DNA during replication. After subtracting any turns that have been added by rotation of the cylinder, the remaining turns can be converted to the location of the fork by multiplying by the helical pitch of DNA (10.5 bp/turn).

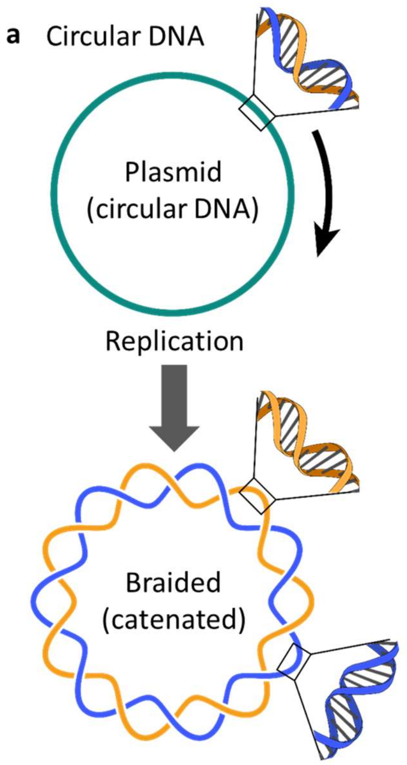


**Fig. S1.** Cartoons explaining that replication generates (+) torsion.

*In vivo*, topoisomerases are essential for relaxation of torsional stress. To illustrate how torsion can accumulate, we consider a special case – the DNA topology in the absence of topoisomerases.

**a**. Circular DNA. Upon replication, one DNA molecule is duplicated into two. The two molecules must be so extensively intertwined (1 catenation per 10.5 bp) that this configuration would be physically impossible due to steric hindrance of the two molecules.

**b**. Linear DNA. The extra turns generated by replication can in principle be dissipated at the ends of a linear DNA. However, *in vivo*, DNA ends are thought to be confined by topological boundaries and barriers that prevent complete supercoiling dissipation. Thus, the two daughter stands are also intertwined.

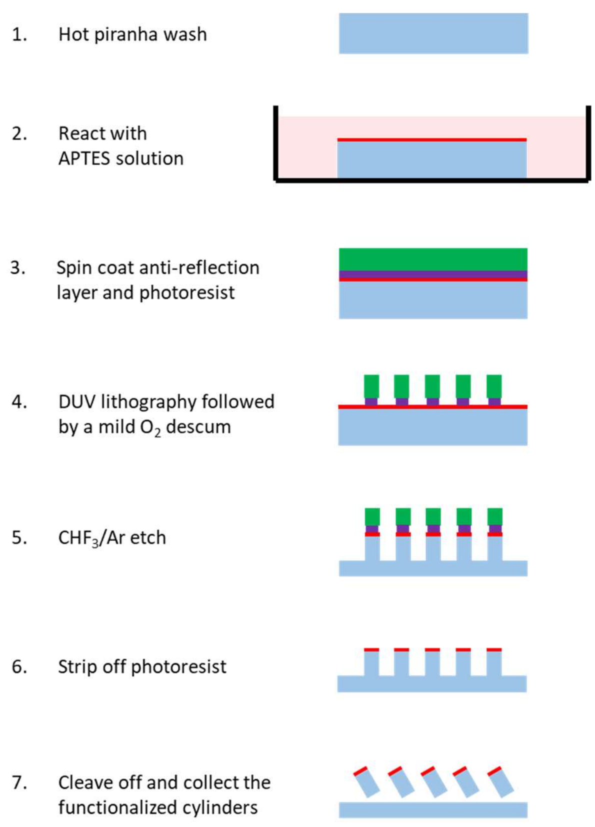


**Fig. S2.** Quartz cylinder fabrication.

This protocol is revised from our previous methods(*6, 21*). 1 and 2, the x-cut single-crystal quartz wafer is cleaned with a hot piranha wash and derivatized by reaction with 3-aminopropyltriethoxysilane (APTES) solution (red). 3, the wafer is spin-coated with an anti-reflection layer (purple) and photoresist (green). 4, the photoresist is exposed with deep UV (DUV) photolithography followed by a mild oxygen ($O_2$) plasma descumming treatment. 5, the pillars are etched into the quartz wafer with a $CHF_3$/Ar plasma etch. 6, the photoresist is stripped from the cylinders, exposing the derivatized top surface that will later be used for specific functionalization. 7, the cylinders are mechanically separated from the wafer by scraping with a razor blade.

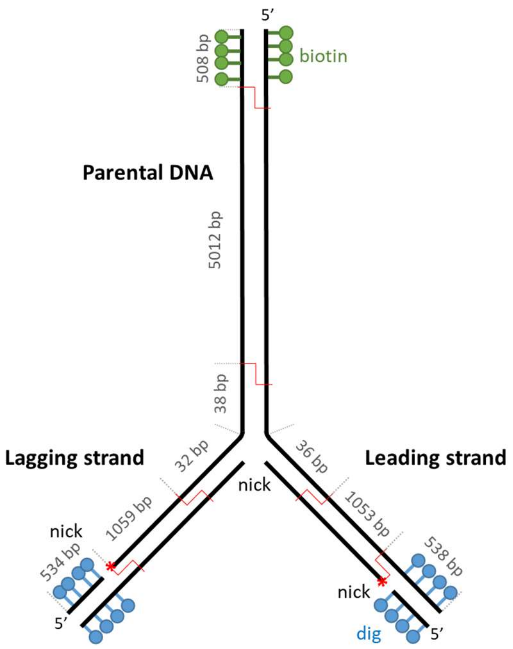


**Fig. S3.** The Y-shaped DNA substrate for replication.

This substrate consists of three segments of DNA brought together at a three-way junction that resembles a replication fork. The parental DNA contains a multi-labeled biotin adapter at the end to enable torsional constraint to a surface of interest. The leading and lagging strands each contain a multi-labeled digoxigenin adapter at the end. Each strand can freely rotate around its own helical axis due to the presence of a nick near the adapter. This nick was introduced to resemble the torsional state *in vivo* since prior studies suggest that each strand can rotate around its own helical axis(*5*).

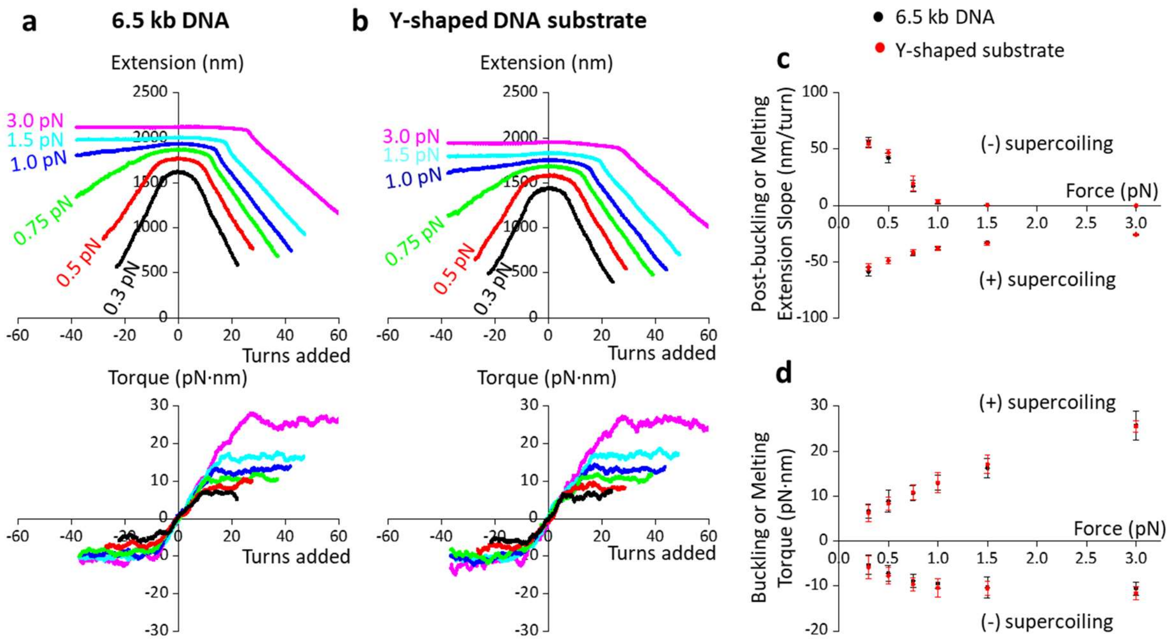


**Fig. S4.** Torsional mechanics of the Y-shaped DNA substrate.

We compared the torsional mechanics of the Y-shaped DNA substrate (Fig. S3) and a linear DNA of similar length (6.5 kb DNA) that we previously employed(*7*). As shown below, we found that the two DNA substrates show nearly identical buckling properties, suggesting that the presence of the leading and lagging anchoring strands do not significantly alter these properties.

**a**. Extension and torque versus turns for the 6.5 kb DNA. Data shown are collected from *N* = 16 individual DNA substrates.

**b**. Extension and torque versus turns for the Y-shaped DNA substrate. Data shown are collected from *N* = 15 individual DNA substrates.

**c**. Post-buckling or melting extension slope versus force. At each force, the slope is obtained from a linear fit to the post-buckling or melting region of the extension for each trace with the error bar being SD of the fit parameter.

**d**. Post-buckling or melting torque versus force. At each force, the torque is obtained from a horizontal line fit to the post-buckling or melting region of the torque for each trace with the error bar being SD of the fit parameter.

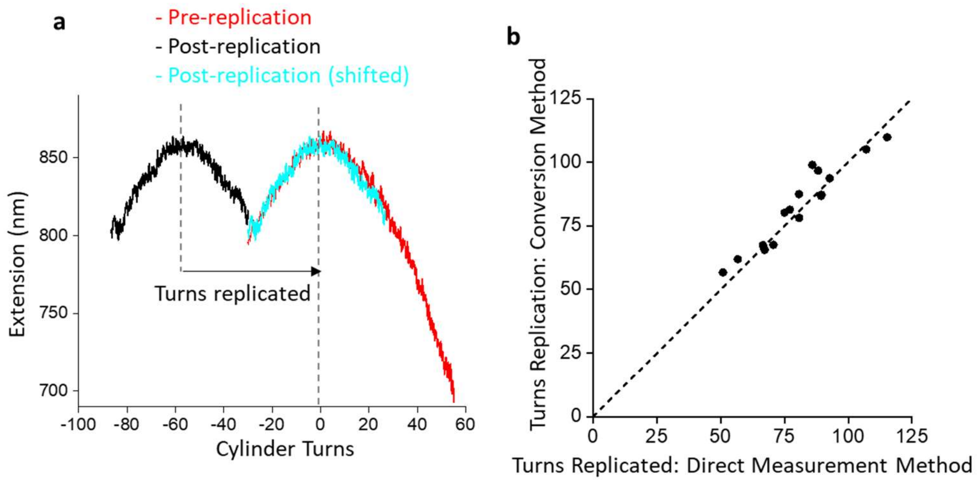


**Fig. S5.** Replisome fork position determination.

During a stalling experiment (Fig. 2), we measured the force and extension of the DNA versus time and used them to determine the fork position versus time (Methods). This conversion method uses a calibration curve to convert measured changes in extension to the replisome fork position. This method assumes that changes in extension are dominated by turns added by the progression of the replication fork. In order to evaluate the accuracy of this conversion method, we developed a direct measurement method by determining the fork position accurately before replisome activity and again at the end of a measurement (**a**). While the conversion method provides the fork position during the measurement, the direct measurement method can only provide the final, stalled replication fork position. We then compare the fork position determined using the conversion method and the direct measurement method (**b**).

**a**. An example trace illustrating the method for direct determination of the fork position at the end of a measurement. In this trace, before replication start, we measured the extension versus turns relation of the Y-shaped DNA substrate under a constant trap height (red). The peak position of this curve represents the equilibrium position of the substrate and was initially centered at zero turns. After replication, we re-measured this relation (black), which was shifted to the left because replication added extra turns to the substrate. The shift in the equilibrium position provides an accurate measure of the number of turns replicated. The conversion to the number of base pairs replicated is straightforward (10.5 bp/turn). Note that this method can only be used for traces that show minimal replication activity during the post-replication measurement as replication activity distorts this relation.

**b**. Comparison of the turns replicated at the end of a stalling measurement using the conversion method and the direct measurement method. Each data point is from a unique trace. Data points falling along the dashed grey guideline (slope = 1) should indicate that the two methods are in close agreement. This plot suggests that the fork position conversion method is accurate within 10%.

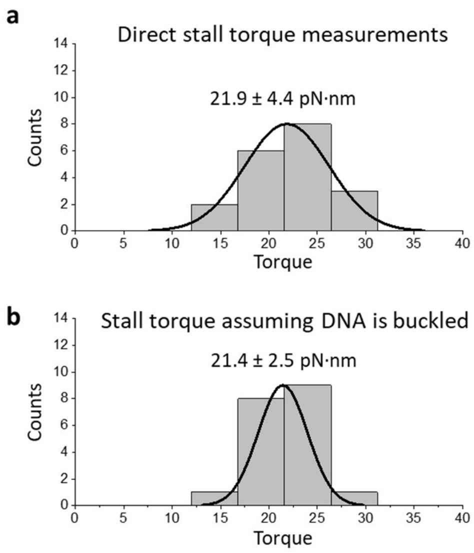


**Fig. S6.** Replisome stall torque measurements.

In Fig. 2, we directly measured the stall torque of the replisome using the torque detector of the AOT. This method is necessary especially when the parental DNA is not buckled. If the parental DNA is buckled, the stall torque can also be obtained from the measured force using the buckling torque versus force relation (Fig. S4d). We have previously used the latter method to determine the torque that RNA polymerase generates during stalling under torsion(*22, 23*). This latter method is advantageous since the noise in force measurements is significantly less than the noise in the torque measurements. Below, we provide evidence that these two methods give similar stall torque values for a replisome, indicating that the parental DNA is buckled when the replisome is stalled. For torque data shown in Fig. 3a and Fig. 4b-d, the latter method is used.

**a.** Stall torque measured directly by the torque detector of the AOT. This histogram is identical to the wt replisome panel shown in Fig. 2. The mean stall torque is 21.9 ± 4.4 pN·nm (mean ± SD) from $N$ = 19 traces.

**b**. Stall torque assuming the parental DNA is buckled. For the same set of traces used in **a**, the stall torque is obtained from the measured force at the stall, converted using the buckling torque versus force relation (Fig. S4). The mean stall torque is 21.4 ± 2.5 pN·nm (mean ± SD).

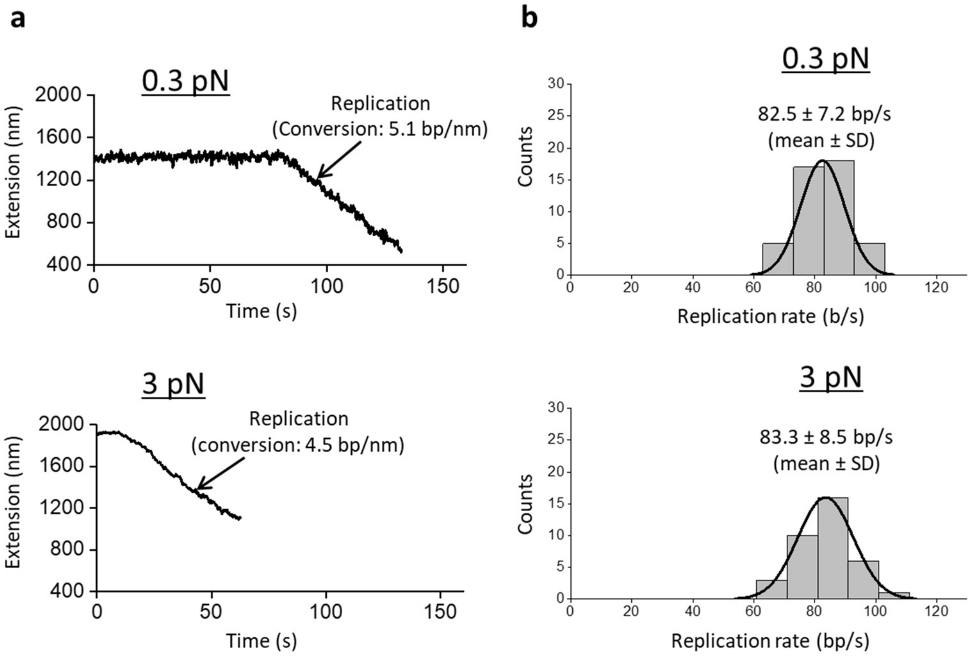


**Fig. S7.** Force on the Y-shaped DNA substrate does not impact replication rate.

During a replisome stalling measurement using the AOT (Figs. 2 and 3), a force is exerted on the Y-shaped DNA substrate in addition to torque and may slow the replication speed. To investigate this possibility, we conducted the following experiments using the MT. In these experiments, we hold a Y-shaped DNA substrate under a specified force. After verifying a DNA tether was torsionally constrained, we introduce to the sample chamber nicking enzyme Nt.BspQI (NEB, Ipswich, MA)that has a target site on the parental DNA. This treatment torsionally un-constrains the tether, and subsequent replication cannot accumulate torsion. We then investigate how the exerted force impacted the replication rate by monitoring the DNA extension.

**a**. Example traces under 0.3 and 3 pN force. Replication decreases the DNA extension since the ssDNA of the lagging strand has a large stiffness(*8, 24*). A trace is terminated if the replication encounters a nicking site. The conversion factor from extension change to base pairs replicated is also indicated.

**b**. Histograms of replication rates under 0.3 and 3 pN force. The mean replication rates are in good agreement with each other, indicating that the applied force does not impact replication significantly.

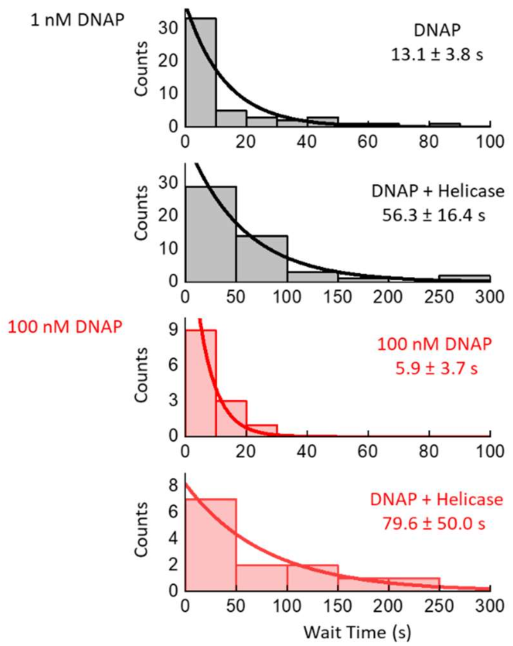


**Fig. S8.** Wait time histograms of DNAP or DNAP+helicase from AOT data.

The reaction buffer contains either the standard protein concentrations (i.e., 1 nM DNAP with other proteins) (Methods) or the standard protein concentrations except for an excess DNAP concentration (100 nM with other proteins). During an experiment, the Y-shaped DNA substrate is unwound using the AOT (Methods). Initial DNAP replication activity is evidenced by an increase in DNA extension as DNAP removes the (-) supercoiling. Subsequent helicase binding (DNAP+helicase) is evidenced by a torque generation of $\geq$ 12.6 pN·nm. The wait times for the initial DNAP activity and the subsequent DNAP+helicase activity are obtained starting from the end of the unwinding step. Each histogram is fit by an exponential function. The mean wait time from the fit and the uncertainty in the fit parameter are shown.

These histograms show that DNAP can locate the fork rapidly (~ 13 s) even at 1 nM concentration. We have also included the wait time histograms for DNAP+helicase. Since DNAP must replicate first to generate a ssDNA region before helicase can load, the combined action of DNAP+helicase requires significantly more time to start.

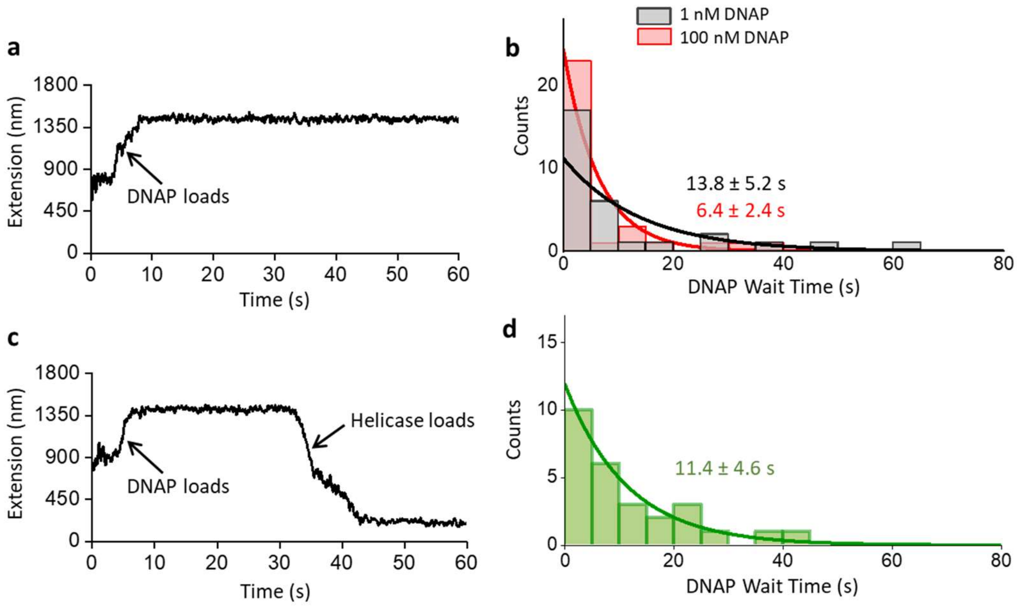


**Fig. S9**. Wait time histograms of DNAP during experiments using the MT.

In these experiments, the Y-shaped template was held under 0.5 pN and was unwound 30 turns by the magnetic tweezers to facilitate replication. Experiments were conducted in the presence of DNAP without or with helicase. The wait time for the start of DNAP replication begins with the end of the initial DNA unwinding step. Only the DNAP wait time is investigated here.

**a**. An example trace in the absence of helicase. DNAP replication activity is evidenced by an increase in the extension when DNAP removes the (-) supercoiling in the parental DNA.

**b**. Histograms of DNAP wait time in the absence of helicase. Shown are the measured histograms and their exponential fits. The numbers are the mean wait times from the fits and the fit uncertainties.

**c**. An example trace in the presence of helicase. DNAP replication activity is evidenced by an increase in the extension when DNAP removes the (-) supercoiling in the parental DNA. Subsequently, helicase activity is evidenced by a decrease in the extension after the arrival of helicase to the fork, (+) supercoiling the parental DNA.

**d**. Histogram of DNAP wait time in the presence of helicase (1 nM DNAP and 180 nM helicase). Shown are the measured histogram and its exponential fit. The numbers are the mean wait time from the fit and the fit uncertainty.

The histograms in **b** and **d** show that DNAP binding to the fork is rapid (~ 10 s) even at 1 nM concentration, and the presence of helicase does not significantly change this binding rate.

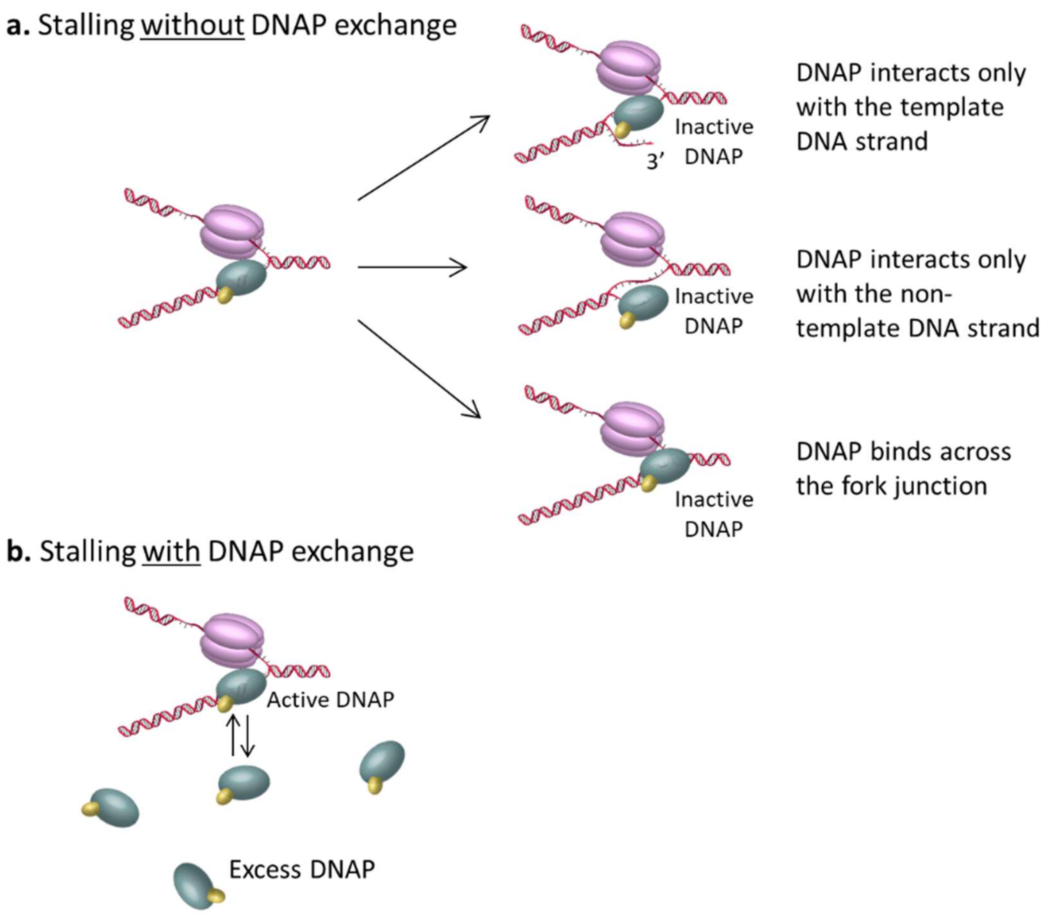


**Fig. S10.** DNA exchange maintains an active fork during stalling by torsion. Our data demonstrate that DNAP exchange during replication stalling under torsion helps maintain an active fork. Without this exchange, the fork is prone to inactivation. The exact cause for this inactivation is unclear. We speculate several inactivated DNAP configurations: DNAP binds to only the template strand DNA, DNAP binds to only the non-template strand DNA, and DNAP binds across the fork junction.

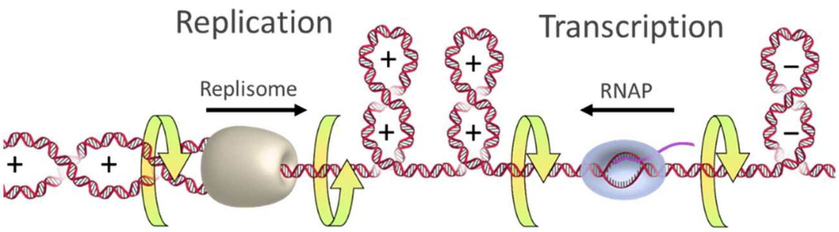


**Fig. S11.** A cartoon illustrating the accumulation of (+) torsion during a replication-transcription head-on conflict.

Both the replisome and RNAP generate (+) supercoiling ahead, accumulating (+) supercoiling or (+) torsion between the two machineries. Since torsion can act over distance, this impact may be experienced by each machinery well before the physical encounter of the two. If the replisome is a more powerful torsional motor, then it can continue to move forward even after torsion stalls the RNAP.

**Movie S1.** Real-time visualization of replisome rotation of DNA.

This video animation shows an example trace of replisome rotation of DNA under 12.6 pN·nm torque as shown in Fig. 1d. The replisome rotation is visualized via the rotation the trapped nanofabricated quartz cylinder in the AOT, as the cylinder rotates to follow the replisome rotation. The inset circle represents the top view of the cylinder with its angular orientation indicated by the red arrow. Only a portion of this trace (from 40 s to 55 s) is animated. As shown, the continuous replication (red regions of the curve) is interrupted by pauses (black regions of the curve). The scale bar provides the conversion to the translocation distance of the replisome.
